# Chloride-dependent conformational changes in the GlyT1 glycine transporter

**DOI:** 10.1101/2020.08.20.259572

**Authors:** Yuan-Wei Zhang, Stacy Uchendu, Vanessa Leone, Richard T. Bradshaw, Ntumba Sangwa, Lucy R. Forrest, Gary Rudnick

**Affiliations:** School of Life Sciences, Guangzhou University, Higher Education Mega Center, 510006 Guangzhou, China; Department of Pharmacology, Yale University School of Medicine, 333 Cedar Street, New Haven, CT 06520-8066; Computational Structural Biology Section, National Institute of Neurological Disorders and Stroke, National Institutes of Health, Bethesda, MD 20892

**Keywords:** Transport, Glycine, Chloride, Mechanism, Structure

## Abstract

The human GlyT1 glycine transporter requires chloride for its function. However, the mechanism by which Cl^-^ exerts its influence is unknown. To examine the role that Cl^-^ plays in the transport cycle, we measured the effect of Cl^-^ on both glycine binding and conformational changes. The ability of glycine to displace the high-affinity radioligand [^3^H]CHIBA-3007 required Na^+^ and was potentiated over 1000-fold by Cl^-^. We generated GlyT1b mutants containing reactive cysteine residues in either the extracellular or cytoplasmic permeation pathways and measured changes in the reactivity of those cysteine residues as indicators of conformational changes in response to ions and substrate. Na^+^ increased accessibility in the extracellular pathway and decreased it in the cytoplasmic pathway, consistent with stabilizing an outward-open conformation as observed in other members of this transporter family. In the presence of Na^+^, both glycine and Cl^-^ independently shifted the conformation of GlyT1b toward an outward-closed conformation. Together, Na^+^, glycine and Cl^-^ stabilized an inward-open conformation of GlyT1b. We then examined whether Cl^-^ acts by interacting with a conserved glutamine to allow formation of an ion pair that stabilizes the closed state of the extracellular pathway. Molecular dynamics simulations of a GlyT1 homologue indicated that this ion pair is formed more frequently as that pathway closes. Mutation of the glutamine blocked the effect of Cl^-^, and substituting it with glutamate or lysine resulted in outward- or inward-facing transporter conformations, respectively. These results provide novel and unexpected insight into the role of Cl^-^ in this family of transporters.

## Introduction

The NSS (Neurotransmitter : Sodium Symporter) family, also known as the SLC6 family of transporters, is responsible for the re-uptake of neurotransmitters after secretion into the synapse. These proteins function at the plasma membrane to take up neurotransmitters and other amino acids and to concentrate them in the cytoplasm. The energetic driving forces powering this accumulation are the electrical potential and ionic concentration gradients across the plasma membrane. An early defining characteristic of these transporters was their requirement for Na^+^ and Cl^-^. Many NSS transporters symport (co-transport) both Na^+^ and Cl^-^ with their substrates (1-12), coupling the membrane potential and ion gradients to substrate accumulation (13-16). The Cl^-^ gradient across mammalian plasma membranes, regulated by the activity of Cl^-^-cation symporters (17), can provide either a positive or a negative driving force for Cl^-^-substrate symport. However, the dependence on Cl^-^ is not universal: prokaryotic NSS transporters, as well as several NSS transporters in animals, do not require Cl^-^ (18-22).

The mechanism of NSS transporter action involves two key conformational changes involving rearrangement of a bundle domain within a scaffold comprising the rest of the structure (23, 24). These rearrangements, observed in several NSS transporters (25-30), open and close aqueous permeation pathways that connect the central substrate binding site alternately with the cytoplasm and extracellular medium. The first conformational change occurs when Na^+^ binds to a site (Na2) (19) that stabilizes a closed cytoplasmic pathway, allowing the extracellular pathway to open (25, 26, 31, 32). The second change occurs when substrate binds, stabilizing specific interactions between the scaffold and bundle domains that involve a Na^+^ ion at the Na1 site, according to results with prokaryotic NSS amino acid transporters that do not require Cl^-^ (28). These interactions, together with additional interactions in the extracellular pathway, overcome the effect of Na^+^ at Na2 and act to stabilize the closed extracellular pathway, allowing the cytoplasmic pathway to open and substrates to dissociate (25, 27, 28). For Cl^-^-dependent NSS transporters, the mechanism by which Cl^-^ participates in such conformational changes, if indeed it does, is unknown.

The site of Cl^-^ binding was first identified by computational analysis and mutagenesis. Comparing the sequence of LeuT, a bacterial NSS amino acid transporter that does not require Cl^-^, with sequences of Cl^-^-dependent mammalian neurotransmitter transporters led two groups to conclude that the position of the ionized γ-carboxyl group of Glu290 in LeuT (Fig. 1) corresponds to the Cl^-^ binding site in Cl^-^-dependent transporters (33, 34). In the Cl^-^-dependent serotonin transporter (SERT), dopamine transporter (DAT) and γ-aminobutyric acid (GABA) transporter (GAT-1), the residue corresponding to LeuT Glu290 was found to be a serine (Ser 339 in GlyT1). Mutating this serine residue in GAT-1, SERT or DAT to either glutamate or aspartate allowed substrate transport in the absence of Cl^-^ (33, 34). Furthermore, replacing Glu290 in LeuT, or the corresponding aspartate in TnaT (another bacterial Cl^-^-independent NSS transporter), with serine imposed a Cl^-^ requirement for substrate binding (34) or transport (35).

**Figure 1.**
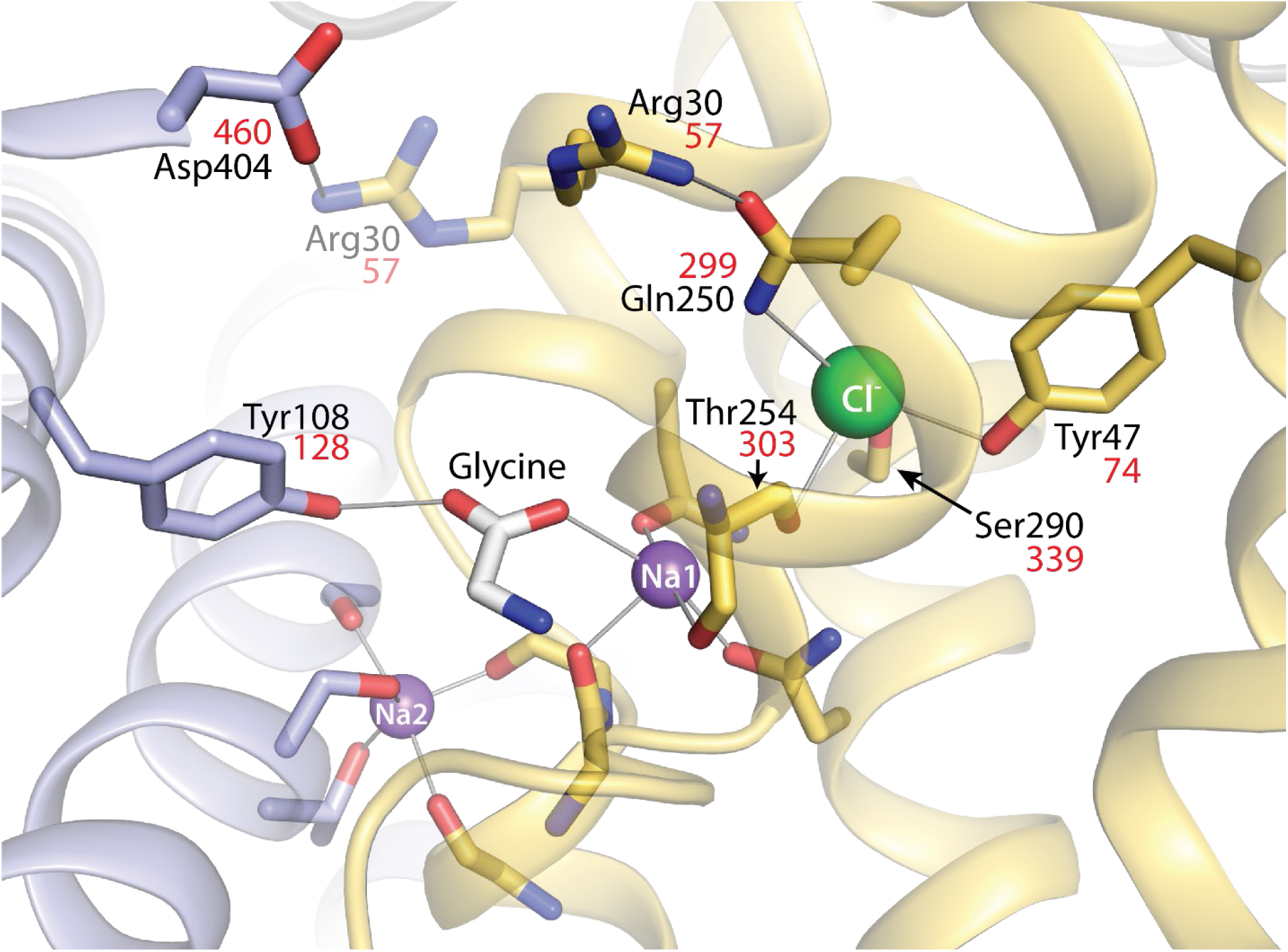
LeuT-E290S with bound Cl^-^, showing interaction networks proposed to account for the effect of ligand binding on conformational change. Outward-occluded conformation of LeuT E290S, a Cl^-^-dependent mutant ((36); PDB 4HOD). Bundle helices are in gold and scaffold helices in light blue, and key residues are shown as sticks. A leucine molecule in the substrate site was mutated to glycine and is shown as sticks with the C atoms in white. Ions are shown as spheres for sodium (purple) and chloride (green). Two ligand-dependent networks of interaction connect the scaffold and bundle domains across the extracellular pathway, as indicated by thin black lines. One of these pathways involves the carboxyl group of substrate connecting Tyr108 from the scaffold domain and Na1 in the bundle domain (28). The other network involves an ion pair between Asp404 (scaffold) and Arg30 (bundle). Arg30 is shown with two rotamers, one (from the LeuT E290S structure) interacting with Gln250 in the Cl^-^ binding site and the other (semi-transparent) interacting with Asp404. For clarity, only residues participating in the substrate- and Cl^-^-dependent conformational changes and the Cl^-^ binding site are numbered. Na^+^coordinating residues (not labeled) include Ala22, Asn27, Thr254, and Asn286 (Na1) and Gly20, Val23, Ala351, Thr354 and Ser355 (Na2) (19). Residue numbers in red are the corresponding positions in GlyT1b (43).

The X-ray structures of LeuT E290S, dopamine transporter (DAT), and SERT (36-38) confirmed the location of the Cl^-^ binding site, near the Na1 site, as shown in Figure 1 for LeuT. The structures of SERT and DAT also confirmed that the coordination of Cl^-^ includes the serine residue corresponding to Glu290 in LeuT (Fig. 1). Additionally, a conserved glutamine residue (Gln250 in LeuT, Fig. 1) was shown to coordinate the bound Cl^-^ ion, which is consistent with the fact that mutation of the corresponding glutamine in GAT-1 affected transport activity and its response to Cl^-^ (39).

To understand the mechanism by which Cl^-^ participates in transport within the NSS family, we focus here on a glycine transporter, GlyT1, which has been shown to symport glycine together with two Na^+^ ions and one Cl^-^ ion in a tightly coupled reaction (40). Both LeuT and GlyT1 transport glycine, suggesting that substrate and Na^+^ interactions are similar in the two proteins, but GlyT1 additionally transports Cl^-^, making it an excellent model system for addressing the Cl^-^-dependence of NSS transporters.

We present here a mechanism by which substrate and Cl^-^ can act independently to induce significant changes in the extracellular pathway of GlyT1. However, only when substrate and Cl^-^ are present together, and in the presence of Na^+^, does this pathway close and the cytoplasmic pathway open, thereby enabling translocation of Na^+^, Cl^-^ and glycine to the cytoplasm.

## Results

### Ion dependence of ligand and substrate binding by GlyT1b

To understand the ionic requirements for substrate binding, and as an aid to following conformational changes in GlyT1b, we used [^3^H]CHIBA-3007 (structure in Fig. 2B inset), a high-affinity GlyT1-selective radioligand (41). We measured its binding to membranes from WT GlyT1b-expressing HEK MSR cells using a filtration assay. Figure 2A shows the binding of tracer [^3^H]CHIBA-3007 (1 nM), its displacement by unlabeled ligand and its dependence on Na^+^ and Cl^-^. In the presence of both ions (open squares) CHIBA-3007 affinity was maximal (K_D_ = 4.85 nM), but even in the absence of either ion, binding was robust, with a K_D_ of 140 nM (filled circles). Because approximately 20% of the 1 nM [^3^H]CHIBA-3007 binding measured in NaCl remained in the absence of these ions (Fig. 2A insert), we used the ability of glycine to displace this radioligand to estimate the ion dependence of substrate binding. Figure 2B shows that the highest glycine affinity (K_I_ = 330 μM) was observed in NaCl (open squares). In the absence of Na^+^, glycine binding was minimal, and was not enhanced by Cl^-^. Although we observed some displacement of [^3^H]CHIBA-3007 binding by glycine in the presence of Na^+^ alone (filled squares), it was weak and incomplete. However, in the presence of Na^+^, Cl^-^ increased glycine affinity over 1000-fold.

**Figure 2.**
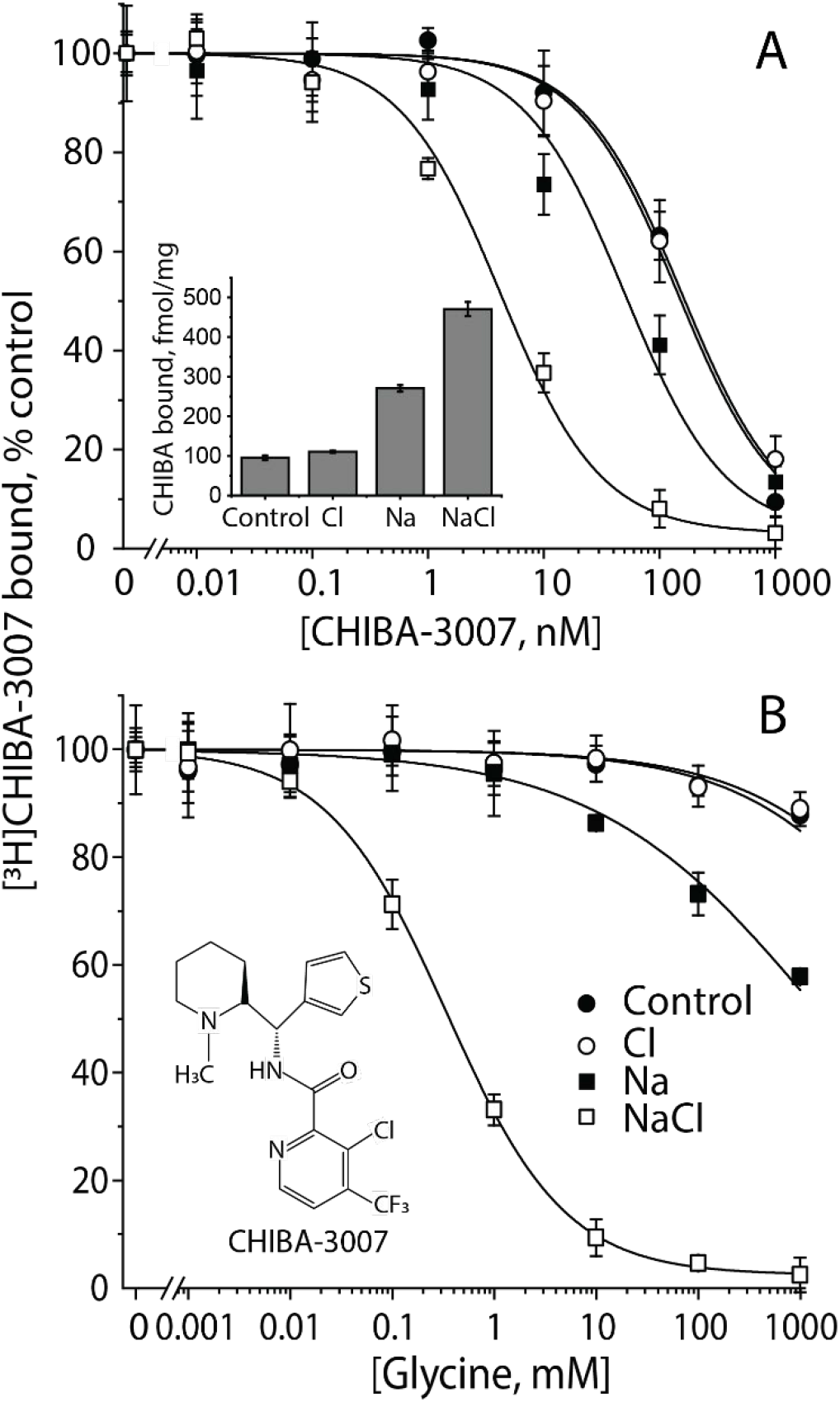
Characterization of ^3^H-CHIBA-3007. **A.** Ion dependence of [^3^H]CHIBA-3007 binding. CHIBA binding was measured by incubating membrane fractions prepared from the cells expressing WT hGlyT1b with [^3^H]CHIBA-3007 in binding buffer with or without Na^+^, Cl^-^, or both. Equimolar NMDG^+^and gluconate were used to replace Na^+^ and Cl^-^, respectively. In experiments measuring displacement of [^3^H]CHIBA-3007 with unlabeled CHIBA-3007, membranes were incubated with radiolabeled CHIBA-3007 (kept at 1 nM) together with unlabeled CHIBA-3007 at 0-1000 nM. The graph shows a representative experiment, with binding expressed as a percentage of that measured in the absence of unlabeled CHIBA-3007. All error bars shown in the figure represent standard deviations from triplicate measurements. The K_D_ values for CHIBA-3007 were estimated to be 140.92 ± 1.77 nM in the absence of Na^+^ and Cl^-^ (control), 141.26 ± 9.89 nM (Cl^-^ alone), 41.98 ± 5.58 (Na^+^ alone), and 4.85 ± 0.45 nM (NaCl), respectively. These calculated values represent the mean ± SEM of three experiments with triplicate measurements. *Inset*, ion dependence of 1 nM [^3^H]CHIBA-3007 binding to WT GlyT1b membranes. Binding, expressed per mg of membrane protein, was corrected for nonspecific binding using membranes from untransfected cells. Error bars represent standard deviations from triplicate measurements. B. Glycine displacement of CHIBA-3007 binding. Glycine was added at final concentrations from 0 -1000 mM along with [3H]CHIBA-3007 (1 nM). The graph shows a representative experiment. All error bars shown in the figure represent standard deviations from triplicate measurements. At concentrations up to 1 M, glycine displaced less than 5% of CHIBA-3007 binding in the absence of Na+. The estimated KI values for displacement by glycine were >1M with Na+ alone and 0.33 ± 0.01 mM in NaCl. This calculated value represents the mean ± SEM of three experiments, each with triplicate measurements.

### Cl^-^-dependence of glycine transport by GlyT1b

Table 1 shows that, as demonstrated previously (3, 42, 43), transport by wild type GlyT1b is dependent on Cl^-^, consistent with Cl^-^-glycine symport (40). To test whether the Cl^-^ binding site in GlyT1b is in the same location as in other neurotransmitter transporters, we examined the role of the conserved serine that has been shown to coordinate Cl^-^ in the neurotransmitter transporters SERT and GAT1, as mentioned above (33, 34, 36-38, 44). In the Cl^-^-independent prokaryotic NSS transporter LeuT, the corresponding residue is a glutamate at position 290 (19, 36), and in GlyT1b it is Ser339 (43). As in GAT1, SERT and DAT (33, 34), replacing Ser339 with aspartate or glutamate led to a loss of Cl^-^-dependence. When normalized for decreased surface expression, the transport activities of S339D and S339E were 68% and 44%, respectively, of wild type activity (Table 1). However, the presence of a carboxylate-containing side chain at position 339 in these mutants largely removed the requirement for Cl^-^, with over 80% of activity remaining in the absence of Cl^-^. By comparison, wild type GlyT1b retained approximately 6%of its activity in the absence of Cl^-^ (Table 1). These results support the expectation that the GlyT1b Cl^-^ site is similar to the Cl^-^ sites characterized in other neurotransmitter transporters.

**Table 1.**
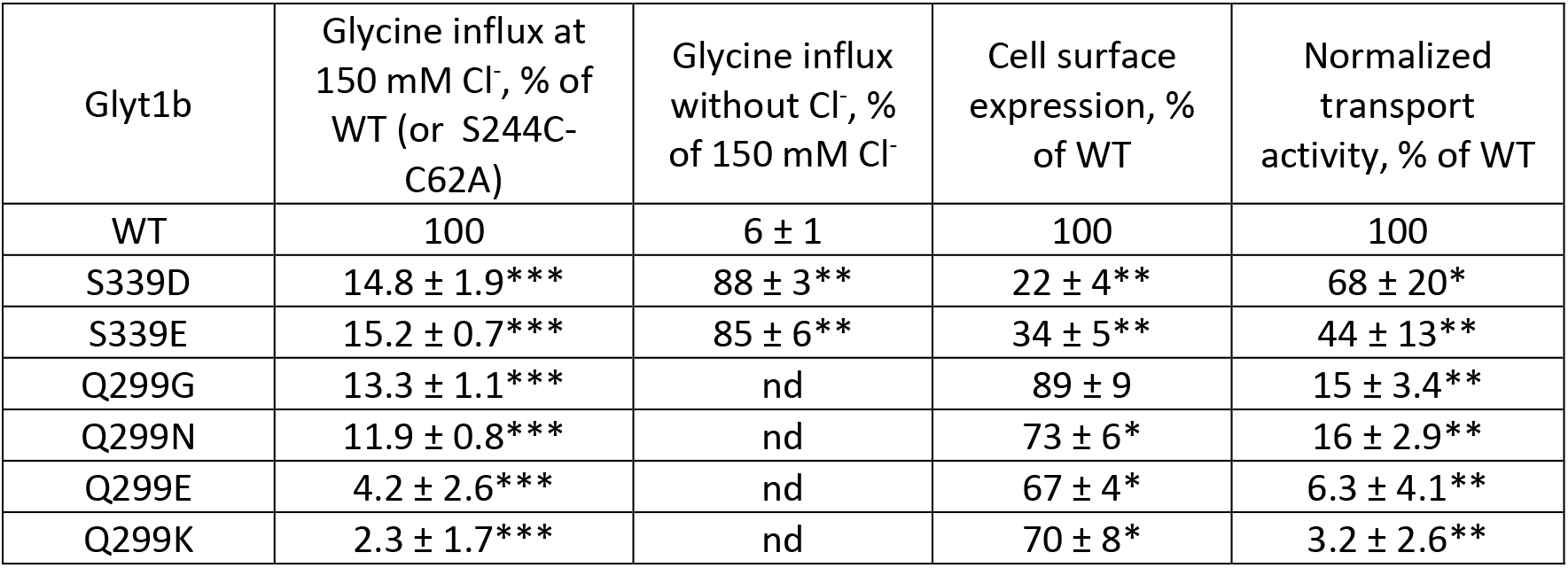
Activity and expression of GlyT1b mutations at Ser399 and Gln299. For WT and Ser339 mutants, [^3^H]glycine (50 nM) influx was measured in both HBS buffer containing 150 mM NaCl and Na-gluconate as described under “Methods”. Activity and surface expression levels for Gln299 mutants are relative to the S244C-C62A background construct, in which Gln299 mutants were generated. Cell surface expression was determined by biotinylation as described under “Methods”. Normalized transport activity shows the influx rate corrected for the surface expression level, relative to WT or S244C-C62A. The values represent means ± SEM; n = 3 for glycine influx; n = 4 for cell surface expression. Asterisks indicate significant differences from WT (* P< 0.05, ** P< 0.01, *** P<0.001) using Student’s t test. nd, not determined.

| Glyt1b | Glycine influx at 150 mM Cl<sup>-</sup>, % of WT (or S244C-C62A) | Glycine influx without Cl<sup>-</sup>, % of 150 mM Cl<sup>-</sup> | Cell surface expression, % of WT | Normalized transport activity, % of WT |
| --- | --- | --- | --- | --- |
| WT | 100 | 6 ± 1 | 100 | 100 |
| S339D | 14.8 ± 1.9*** | 88 ± 3** | 22 ± 4** | 68 ± 20* |
| S339E | 15.2 ± 0.7*** | 85 ± 6** | 34 ± 5** | 44 ± 13** |
| Q299G | 13.3 ± 1.1*** | nd | 89 ± 9 | 15 ± 3.4** |
| Q299N | 11.9 ± 0.8*** | nd | 73 ± 6* | 16 ± 2.9** |
| Q299E | 4.2 ± 2.6*** | nd | 67 ± 4* | 6.3 ± 4.1** |
| Q299K | 2.3 ± 1.7*** | nd | 70 ± 8* | 3.2 ± 2.6** |

### Constructs for measuring conformational changes in GlyT1b

Cys62, near the end of TM1 in GlyT1b, corresponds to a reactive cysteine common to many mammalian NSS transporters. As shown in Table 2, replacing Cys62 with an alanine had little effect on transport, but markedly decreased the rate of inactivation by 2-aminoethyl methane thiosulfonate hydrobromide (MTSEA) and [2-(trimethylammonium)ethyl] methane thiosulfonate bromide (MTSET). These reagents react with cysteine sulfhydryl groups, forming a mixed disulfide with the thioethylamine or thioethyltrimethylammonium moiety, respectively, of the reagent. In the case of wild-type GlyT1b, this reaction dramatically inhibits transport activity and CHIBA-3007 binding. The C62A mutation thus provides a low-reactivity background in which to measure reactivity of cysteine residues inserted at other locations. We chose to use MTSET for subsequent studies because of its lower reactivity with the remaining endogenous cysteine residues.

**Table 2.** Characteristics of Glyt1b background constructs. Transport rates were measured over a range (0.05 –200 μM) of glycine concentrations and surface expression was determined by biotinylation as described under “Methods”. K_M_ and V_max_ values represent the means ± SEM of three experiments for kinetic analysis and four experiments for biotinylation. Sensitivity to MTSEA and MTSET modification was determined from the concentration of MTS reagents and the half-time for inactivation of CHIBA-3007 binding (S244C-C62A) or glycine transport (Y60C-C62A) as described under “Methods”. Asterisks indicate significant differences from WT (* P< 0.01, ** P<0.005) using Student’s t test. nd, not determined.

| Glyt1b | $K_M$ (μM) | $V_{max}$<br>(pmol/min/<br>mg) | Cell surface<br>expression<br>(% of WT) | Rate of<br>binding<br>inactivation<br>by MTSEA,<br>$M^{-1} \cdot sec^{-1}$ | Rate of<br>binding<br>inactivation<br>by MTSET,<br>$M^{-1} \cdot sec^{-1}$ | Rate of<br>transport<br>inactivation<br>by MTSET,<br>$M^{-1} \cdot sec^{-1}$ |
| --- | --- | --- | --- | --- | --- | --- |
| WT | $35.2 \pm 2.1$ | $1560 \pm 93$ | 100 | $3.29 \pm 0.09$ | $1.79 \pm 0.07$ | $0.61 \pm 0.03$ |
| C62A | $36.4 \pm 2.6$ | $1466 \pm 81$ | $89.3 \pm 5.3$ | $0.40 \pm 0.01$ | $0.12 \pm 0.01$ | $0.02 \pm 0.01$ |
| Y60C-C62A | $41 \pm 4$ | $715 \pm 50^{**}$ | $65.3 \pm 4.7^*$ | nd | nd | $6.52 \pm 0.42$ |
| S244C-C62A | $35.2 \pm 2.8$ | $645 \pm 18^{**}$ | $50.7 \pm 5.9^*$ | $32.10 \pm 1.49$ | $20.8 \pm 0.70$ | nd |

To follow conformational changes in GlyT1b, we identified positions in the extracellular and cytoplasmic pathways that, according to accessibility measurements with SERT and LeuT (28, 45), should be more accessible in outward- and inward-open conformations, respectively. According to a structural model of GlyT1 built based on an outward-open *Drosophila* DAT X-ray structure (PDB ID 4XP4) (46) (Fig. S1), Cys60 is exposed to the open extracellular pathway and Cys244 is buried in the interior of the protein (Fig. 3A). By contrast, in a recently-reported structure of GlyT1 in an inward-open conformation (44), Cys60 is buried, due to closure of the extracellular pathway (Fig. 3B). The region containing Cys244 was not resolved in the X-ray structure (dashed line), but was resolved as an extended loop in an inward-open cryo-EM structure of SERT (47). Using GlyT1b C62A, we replaced Tyr60 in TM1 or Ser244 in TM5 with cysteine to measure accessibility of the extracellular and cytoplasmic pathways, respectively, using methods established with SERT (45, 48). The K_M_ values for glycine in these three mutants were similar to those of wild type GlyT1b. V_max_ values for C62A-Y60C and C62A-S244C were about half those of wild type, largely due to decreased surface expression (Table 2).

**Figure 3.**
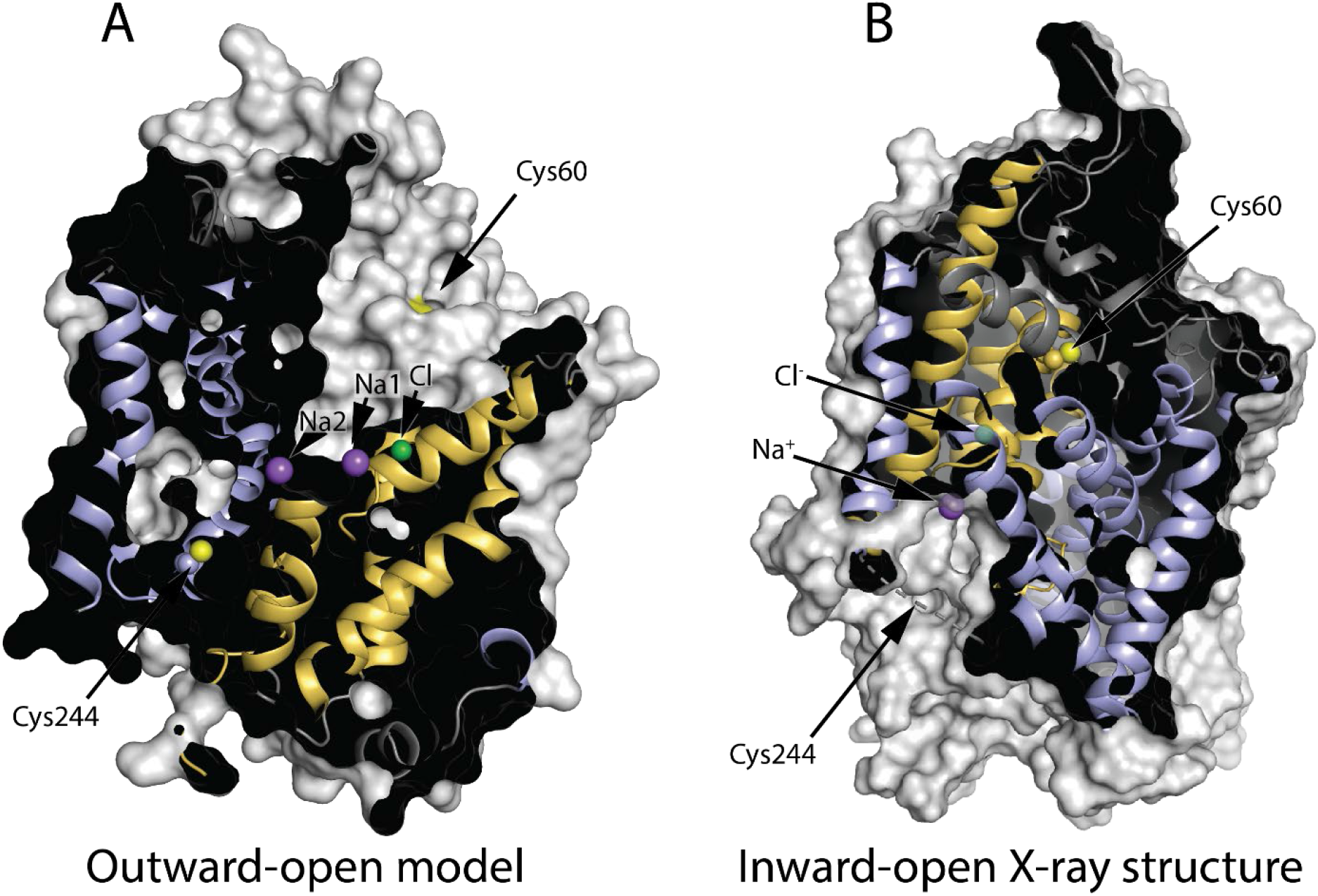
GlyT1 in outward-open (model) and inward-open (X-ray structure) conformations. **A.** In this outward-open model (Fig. S1) showing cysteine at both Tyr60 and Ser244, Cys60 is accessible through the extracellular pathway, but Cys244 is buried. **B.** In the inward-open conformation (PDB ID 6ZPL with the ligand removed) (44), the closed extracellular pathway buries Cys60 within the protein structure Cys244 is in an unresolved region, shown by the dashed line, which is expected to be accessible to the cytoplasm. Residue numbering is based on the published sequence of GlyT1b (43). Helices from the bundle domain are shown in gold and those from the scaffold domain are light blue. Positions for the ion binding sites were superimposed on the model and ions found in the inward-open structure are superimposed in semi-transparent form.

### Na^+^, Cl^-^ and glycine influence GlyT1b conformation

Figure 4A and B show the effect of Na^+^, Cl^-^ and glycine on the reactivity of Cys60 and Cys244. In the absence of Na^+^, Cl^-^ had little effect on transporter conformation. Na^+^, however, increased the reactivity of Cys60 and decreased Cys244 reactivity, consistent with its effect on other NSS transporters (Fig. 4A, B, 3^rd^ column from left). These results suggest that, as observed for LeuT, Tyt1, SERT and DAT, Na^+^ binding at Na2 stabilized outward-open conformations of GlyT1b in which the extracellular pathway is open and the cytoplasmic pathway is closed (25, 26, 28, 31). Addition of Cl^-^ plus glycine reversed the effect of Na^+^ and decreased Cys60 reactivity while it increased Cys244 reactivity, consistent with closing the extracellular pathway and opening the cytoplasmic pathway (rightmost column in Fig. 4A, B). In LeuT and Tyt1, bacterial transporters that do not require Cl^-^, substrate addition similarly reversed the conformational effect of Na^+^ in the absence of Cl^-^ (25-28, 31). In SERT, which does require Cl^-^, the substrate, serotonin, also reversed the Na^+^ effect in the presence of Cl^-^ (30).

**Figure 4.**
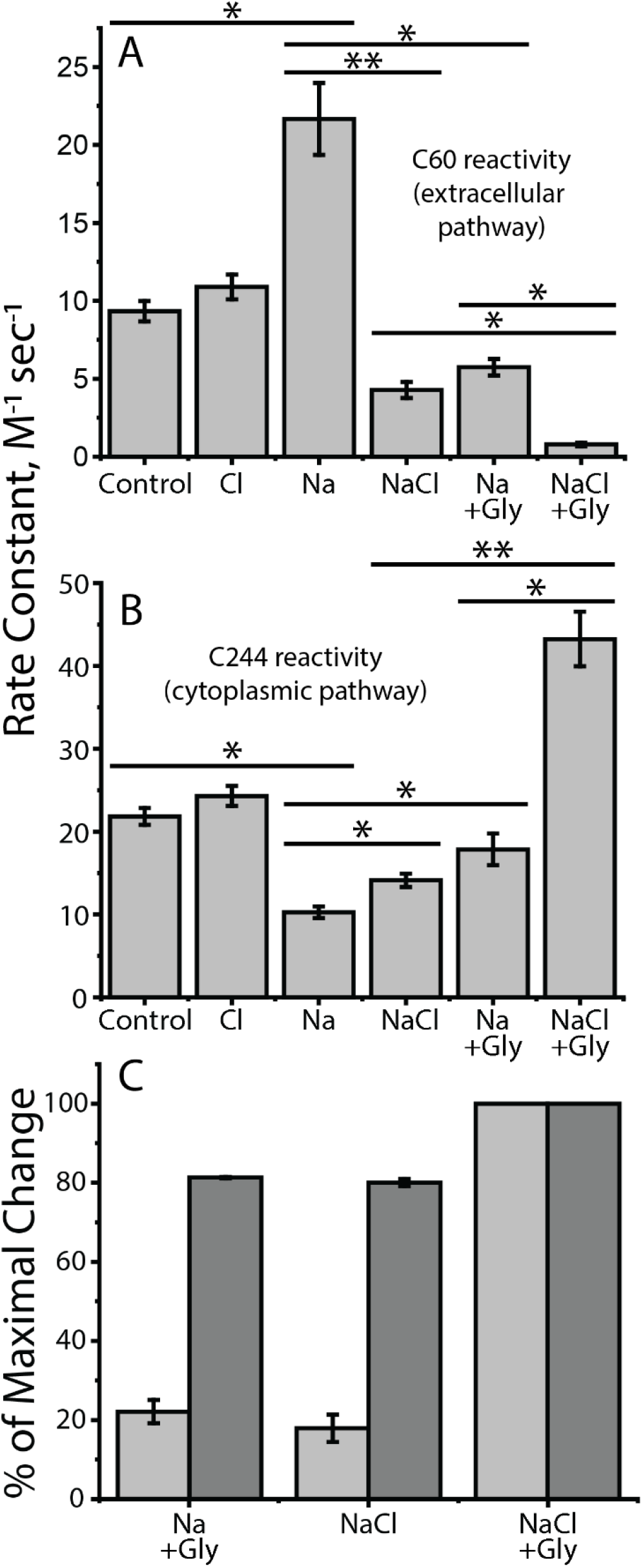
Accessibility changes induced by ions and substrate (glycine). The conformational response to ligand binding was estimated by changes in the reactivity of Cys60 in the extracellular pathway and Cys244 in the cytoplasmic pathway towards MTSET added to the medium of intact cells expressing GlyT1b Y60C-C62A (panel A) or to membranes from cells expressing GlyT1b S244C-C62A (panel B). **A.** Cys60 reactivity; **B.** Cys244 reactivity. Asterisks indicate significantly different rate constants (*, P<0.02; **, P<0.01) using Student’s t test. All error bars represent the SEM; n ≥ 4. **C.**Conformational response, in the presence of Na^+^, to Cl^-^ or glycine, relative to the response to both. The change in accessibility of Cys60 and Cys244 between Na^+^ alone and NaCl + glycine was set at 100% and the changes induced by Cl^-^ or glycine scaled accordingly. Light gray bars represent cytoplasmic pathway (Cys244) reactivity and dark gray bars, that of the extracellular pathway (Cys60). All error bars represent the SEM; n ≥ 4.

In GlyT1b, addition of either Cl^-^ or glycine, in the presence of Na^+^, caused marked changes in accessibility (Fig. 4A, B, 4^th^ and 5^th^ columns from left). These changes were in the opposite direction to the Na^+^-induced stabilization of the outward-open conformation, but not as large as the change due to addition of both glycine and Cl^-^. These results suggest that, in the presence of Na^+^, substrate and Cl^-^ each act independently to favor partial closure of the extracellular pathway and partial opening of the cytoplasmic pathway. Moreover, under these conditions, glycine and Cl^-^ act synergistically, especially on the cytoplasmic pathway, where the changes in response to either agent alone were dramatically amplified by the presence of the other (Fig. 4B). For the small changes observed in the cytoplasmic pathway, our methods do not distinguish between limited conformational changes that increase Cys244 reactivity without fully opening the pathway and full opening of a small fraction of transporters. However, full opening in the absence of all substrates would impair ion-substrate coupling and allow substrate-independent ion flux, neither of which was observed in electrophysiological studies (40).

Our results showed a clear difference between changes in the reactivity of Cys60 when compared with that of Cys244. Figure 4C shows the response, in the presence of Na^+^, to addition of glycine and Cl^-^ as a percentage of the maximal change seen with both. The reactivity of Cys244 in the cytoplasmic pathway increased by only 15-20% of this maximum in the presence of either glycine or Cl^-^ (light gray). In contrast, the reactivity of Cys60 in the extracellular pathway decreased by about 80% of this maximum under the same conditions (dark gray).

### The role of Gln299 in conformational responses to Cl^-^

The ability of Cl^-^ to mediate conformational changes in GlyT1 must depend on specific interactions that form or break in response to ion binding. Here we consider the contributions of the highly conserved glutamine at position 299 in GlyT1b (Fig. 1), which has been invoked as a critical Cl^-^ binding site residue in GAT-1 (34). The role of this position has also been explored in the E290S mutant of LeuT, which requires Cl^-^ for substrate binding (34) and thus serves a model for the Cl^-^-dependent NSS transporters (36). In the X-ray structure of LeuT-E290S, which is in an outward-occluded conformation, Gln250 (corresponding to Gln299 in GlyT1) interacts simultanteously with both the bound Cl^-^ ion and with Arg30 (Arg57 in GlyT1b) from TM1 (36)(Fig. 1). Arg30, in turn, is found to form a salt-bridge with Asp404 (Asp460 in GlyT1b) across the extracellular pathway in a structure of the inward-facing conformation of LeuT (24), and thus may stabilize the closure of this pathway (19, 49). Indeed, even in the outward-occluded conformation represented by LeuT-E290S, i.e., where the extracellular pathway is partially closed and the substrate is occluded by protein side chains, a simple rotamer flip of Arg30 places the side chain within hydrogen-bonding distance of Asp404 (Fig. 1, semi-transparent). Interpretation of the nature of this network using structures alone, however, is challenging, since it appears to be highly dynamic (36). By contrast, molecular dynamics simulations provide a means to interrogate such networks in different states of the conformational cycle. For example, simulations of LeuT perturbed by removal of the negative charge (either by deletion of the Cl^-^ ion in E290S, or by neutralizing the endogenous Glu290 in wild type LeuT) allowed Kantcheva et al. to propose that Cl^-^ binding influences the formation of the Arg30-Asp404 salt bridge through Gln250 (36), at least in outward-open conformations.

To assess whether the interaction network involving Gln250 differs according to the conformational state of the transporter in LeuT, and by extension whether Gln299 might provide a role in the conformational sensing of the chloride ion in GlyT1, we carried out 2 μs-long molecular dynamics simulations of wild-type LeuT based on structures in outward-open (PDB code 3TT1), outward-occluded (PDB code 3F48) and inward-open states (PDB code 3TT3; see Methods). Note that, since the LeuT-E290S structure contains leucine, a poor substrate, in the central binding site, we used a structure of the outward-occluded conformation bound to alanine (PDB code 3F48) for that simulation.

In simulations of the outward-open conformation of LeuT (3TT1) (Fig. 5A), Arg30 interacted transiently with Gln250 (green bar) or with Asp404 (blue bar), or neither (gray bar). Those interactions were anticorrelated, i.e., Gln250 and Asp404 appear to compete for an interaction with Arg30 (Fig. S2A). In this conformation of the protein, Gln250 was also very flexible and not consistently interacting with the negatively-charged Glu290, presumably due to suboptimal packing of the bundle helices, significant hydration of the pathway, or both. In simulations of the outward-occluded conformation, in which the pathway is partially closed, Arg30 formed more persistent salt-bridge interactions with Asp404 (Fig. 5B), and only very occasional forays toward Gln250 (Fig. S2B). In this conformation, Gln250 also interacted more stably with Glu290, apparently due to an optimal arrangement of the helices in the bundle (Fig. S2B).

**Figure 5:**
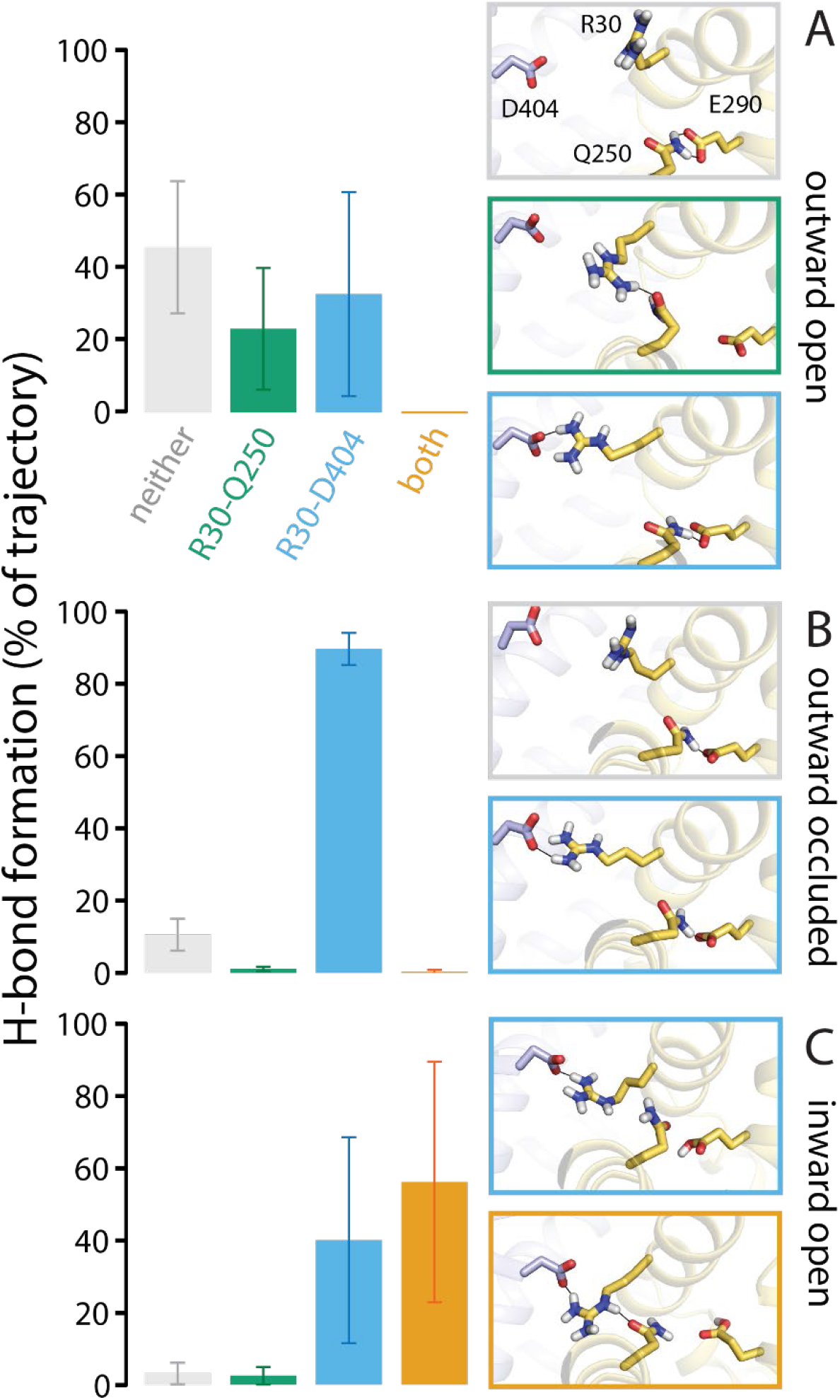
Formation of H-bonds linking the extracellular pathway to the chloride binding site during MD simulations of LeuT. Data represent interactions of R30 exclusively with either Q250 (green bars) or D404 (blue bars), both residues simultaneously (orange bars), or neither (grey bars) for (**A**) outward-open apo (3TT1), (**B**) outward-occluded holo (3F48), and (**C**) inward-open apo (3TT3) conformations of LeuT. Panels indicate snapshots representing the most highly-populated states in each conformational states of the transporter, and the colors of the borders matches that of the bars. An H-bond is counted as present if the minimum distance between the H-bond donor/acceptor atoms of the side-chains is < 3.2 Å, and was computed every 100 ps. The results are averaged over three simulation repeats, each 2 μs long, and the error bars represent the standard deviation.

As mentioned, we also compared the interactions present in simulations of an inward-open conformation of LeuT (PDB code 3TT3). However, it should be noted that the pKa of Glu290 in this conformation is shifted, consistent with this residue having a high probability of being protonated in this state of the transport cycle (50) and thereby conferring proton antiport to LeuT (51). We therefore protonated this residue during the simulations, which to some extent mimics a chloride-free state, and which seemed to slightly destabilize the Gln250-Glu290 interaction (Fig. S2C). Nevertheless, it is clear that the packing of the bundle against the scaffold in this inward-open state allows Arg30 to interact with both Asp404 and Gln250 more consistently than in the outward-facing states (Fig. 5C, S2C). These results support the notion that the interaction network involving Q250 that links the extracellular pathway to Glu290 is strongly conformationally dependent in LeuT. Given that Glu290 is an excellent mimetic of chloride, we infer that the closure of the extracellular pathway in GlyT1 is also likely to be influenced by the presence or absence of the negatively-charged Cl^-^ ion, and as such, these simulations predict that Gln299 will be a key player in that process.

### Gln299 mutants

To test the possibility that Gln299 in GlyT1b controls the interaction between Arg57 and Asp460, we mutated Gln299 to glycine, asparagine, glutamate and lysine in the wild type background. Each of these mutants was expressed well on the cell surface and bound CHIBA-3007, but transported glycine poorly (Table 1). Normalized for surface expression, Q299G and Q299N catalyzed glycine influx at about 15% the rate of wild type but Q299E and Q299K were essentially inactive (Table 1). To assess the nature of this transport defect, we determined the Cl^-^ dependence of glycine transport in Q299G and Q299N (Fig. 6A). In contrast with wild type GlyT1b, which symported Cl^-^ and glycine with a K_M_ for Cl^-^ of 13.5 ± 1.7 mM, transport by Q299G and Q299N increased linearly with Cl^-^ concentrations up to 200 mM, indicating a lower Cl^-^ affinity for these mutants, consistent with Gln299 forming part of the Cl^-^ binding site.

**Figure 6.**
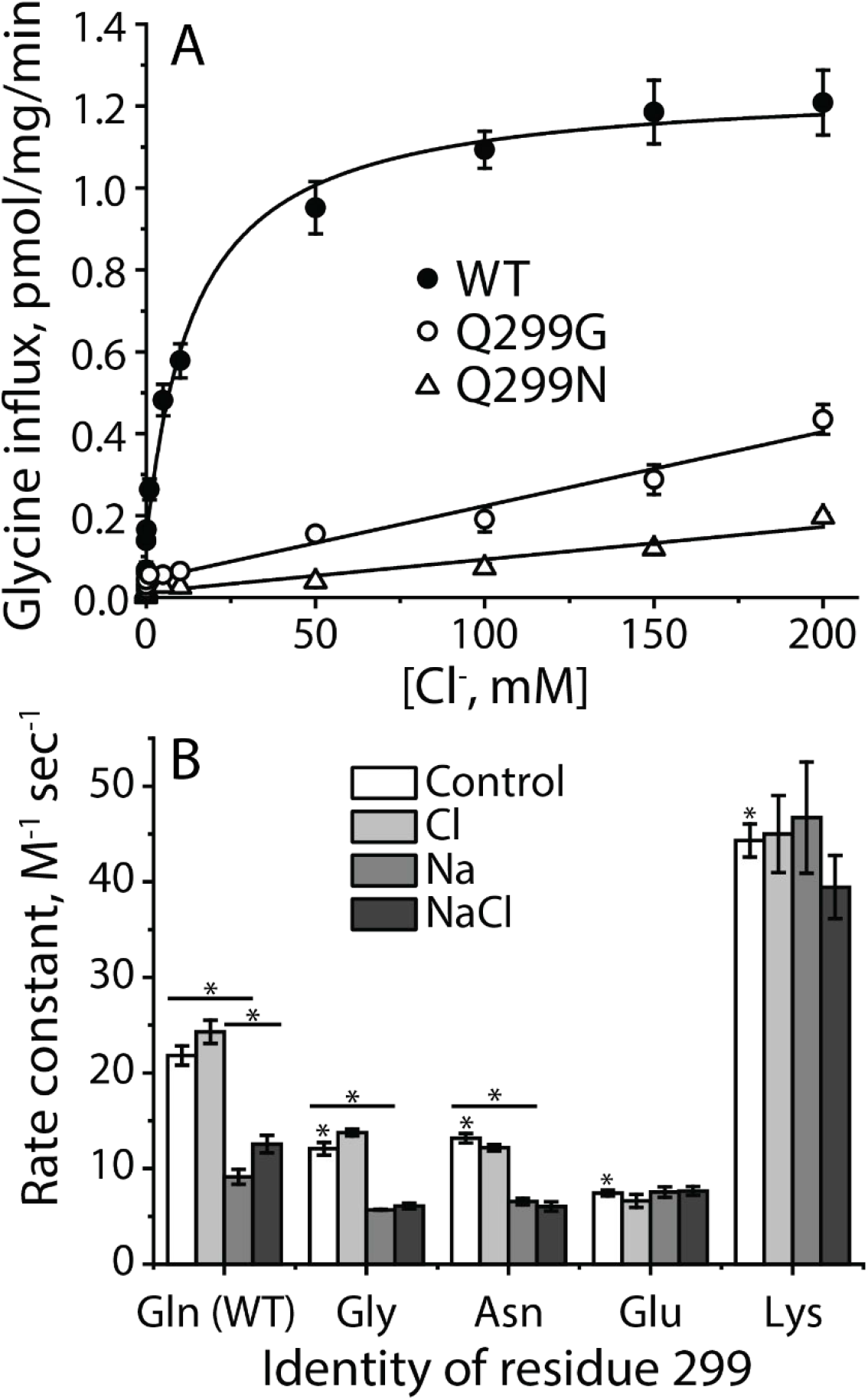
Characterization of Gln-299 mutants. **A.** Chloride dependence of transport. Gluconate was used to replace Cl^-^ for concentrations below 200 mM with [Na^+^] constant at 200 mM. The graph shows a representative experiment. All error bars represent standard deviations from three measurements. The Km value for Cl^-^ with WT was 13.56 ± 1.70 mM. This value represents the mean ± SEM of three experiments. **B.** Gln299 mutants influence GlyT1b conformation and its response to Na^+^ and Cl^-^. Rate constants were determined for MTSET modification of Cys244 in the cytoplasmic pathway of S244C/C62A and Gln299 mutants in the S244C/C62A background. NMDG^+^ and gluconate were used as replacements for Na^+^ and Cl^-^, respectively. Ion concentrations were kept a constant at 150 mM in all experiments, and control represents the absence of NaCl. Control columns with * are significantly different from S244C/C62A, P<0.01. Within each mutant, differences between control and Na^+^, or between Na^+^ and NaCl, where significant, are shown. *, P<0.01, using Student’s t test. All error bars represent the SEM; n ≥ 3.

The ability of the Gln299 mutants to bind CHIBA-3007 allowed us to examine their conformations in the presence of Na^+^ and Cl^-^ even in the absence of transport activity (Fig. 6B). Compared with the background construct GlyT1b C62A-S244C (Fig. 6B, first set of columns), Cys244 in the cytoplasmic pathway was less accessible in Q299G, Q299N and Q299E. As with GlyT1b C62A-S244C, Cl^-^, in the absence of Na^+^, had little effect on Cys244 accessibility (open vs. light gray columns). Na^+^ reduced Cys244 accessibility in Q299G and Q299N as it did in GlyT1b C62A-S244C (Fig. 6B, medium gray columns) but in the presence of Na^+^, the small Cl^-^-dependent increase in Cys244 accessibility characteristic of GlyT1b C62A-S244C was not observed in Q299G and Q299N (Fig. 6B, dark grey columns), possibly due to the low Cl^-^ affinity of these mutants (Fig. 6A).

In the Q299E mutant, the accessibility of Cys244 under control conditions (no Na^+^ or Cl^-^) was lower than for any other mutant – approximately the same as for GlyT1b C62A-S244C, or the Q299G, or Q299N mutants in the presence of Na^+^ (Fig. 6B, fourth set of columns). This accessibility was not affected by Na^+^or Cl^-^. In contrast, the accessibility of Cys244 in GlyT1b C62A-S244C-Q299K was high under control conditions (Fig. 6B, last set of columns) - similar to that of GlyT1b C62A-S244C in the presence of NaCl and glycine (Fig. 4B). As with the Q299E mutant, neither Na^+^ nor Cl^-^ significantly changed this accessibility. These results suggest that a positively or negatively charged side chain at position 299 can drive reactivity of the cysteine replacing Ser244 to opposite extremes consistent with inward- or outward-open GlyT1b conformations.

## Discussion

The results presented here help us to understand the role of Cl^-^ in mammalian SLC6 amino acid transporters. The binding site for Cl^-^ in NSS transporters has been established in functional, computational and structural studies (33-39), but there has been little experimental evidence to demonstrate how Cl^-^ binding is coupled to transport. Our results show that Cl^-^ acts independently of substrate to decrease accessibility in the extracellular pathway and increase it in the cytoplasmic pathway (Fig. 4). We propose that these accessibility changes reflect conformational changes due, at least in part, to Cl^-^ interacting with a conserved glutamine residue (Gln299 in GlyT1b) that, in turn, influences the formation of an ion pair in the extracellular pathway (Figs. 1, 5-6). The ability of Cl^-^ to close the extracellular pathway of GlyT1b was blocked by mutation of Gln299, and the conformation of GlyT1b was dramatically shifted, in opposite directions, by replacement of Gln299 with residues of opposite charges (Fig. 6B).

These findings illustrate a concept of NSS transporters in which two ligand-dependent networks cooperate to close the extracellular pathway and allow the cytoplasmic pathway to open. As we described previously for LeuT (28), one of these networks centers on the interactions of the substrate carboxyl group with a tyrosine in TM3 of the scaffold domain and with Na^+^ bound at the Na1 site in the bundle domain. In LeuT, this residue is Tyr108 (Tyr128 in GlyT1b, Fig. 1) and mutation of Tyr108 blocked the ability of bound substrate to initiate the open-out to open-in conformational change that allows substrate and Na^+^ to dissociate to the cytoplasm (28). The results here show that another network involves formation of an ion pair between the scaffold and bundle domains further towards the extracellular end of the pathway (Arg30-Asp404 in LeuT, Arg57-Asp460 in GlyT1b) (Fig. 1). Formation of this network is facilitated by Cl^-^ binding and depends on Gln299 in GlyT1b (Gln250 in LeuT). Specifically, the interaction between Cl^-^ and Gln299 in GlyT1b apparently disengages Arg57 so that it can interact with Asp460 to close the extracellular pathway (Fig. 1).

Our results show that, even in the absence of substrate, Cl^-^ influences GlyT1b conformation by decreasing accessibility in the extracellular pathway and increasing accessibility in the cytoplasmic pathway (Fig. 4). These changes are consistent with a conformational shift in the transporter toward an inward-open state, and could reflect either that a subset of the transporters have undergone the complete conformational change, or more likely, that the pathways become partially occluded or partially opened, respectively. Likewise, substrate can foster similar accessibility changes in both pathways in the absence of Cl^-^ (Fig. 4). In the extracellular pathway, these changes in accessibility were almost as large as the effect of adding both substrate and Cl^-^ together, but changes in cytoplasmic pathway accessibility were much smaller (Fig. 4C) and therefore not indicative of conversion to a fully inward-open state. It is worth noting that glycine transporters, including GlyT1b, are characterized by tight stoichiometric coupling of Na^+^, substrate and Cl^-^ transport (52). Therefore, we would expect that only when Na^+^, Cl^-^ and substrate are bound would their combined conformational effects close the extracellular pathway and allow the cytoplasmic pathway of GlyT1 to fully open (Fig. 4B) enabling dissociation of ions and substrates. In NaCl or Na^+^+ glycine, GlyT1 may visit (in addition to outward open conformations that allow binding of glycine and Cl^-^, respectively) “inward-occluded” states previously observed for MhsT and LeuT (53, 54) in which the extracellular pathway has closed but the cytoplasmic pathway has not yet opened. These conformations differ from inward-open states by the position of TM1a, which opens the cytoplasmic pathway as it separates from TMs 6 and 8. These structures also show changes in the cytoplasmic pathway, relative to outward-facing states, particularly involving TM5 (where the cysteine replacing Ser244 is located). Such structural changes are consistent with the accessibility changes in both pathways observed here in the presence of Na^+^ and either glycine or Cl^-^, even without both of the latter ligands being present (Fig. 4B).

The possibility that Cl^-^ binding might affect formation of an ion pair between Arg57 and Asp460 (Fig. 1) was first raised in response to mutation of Gln291 in GAT1 (Gln299 in GlyT1b) (39). That paper showed that mutations of Gln291 inhibited transport activity and noted the proximity of Gln291 to Arg69 (Arg57 in GlyT1b), speculating that interaction between Gln291 and Arg69 might be the cause. This interaction was later invoked by Kantcheva et al (36), who used molecular dynamics simulations to predict that a negative charge on Glu290 in LeuT (or the Cl^-^ bound to LeuT-E290S) would affect the formation of the Arg30-Asp404 salt bridge (Arg57-Asp460 in GlyT1b). Their predictions, as well as ours (Fig. 5) identified the possibility that Arg30 in LeuT could interact with either Asp404 or Gln250. Our simulations additionally illustrate the ability of Arg30 to interact simultaneously with both Asp404 and Gln250 in inward-open conformations of LeuT. Although the simulations cannot directly speak to whether Cl^-^binding would enhance or inhibit ion-pair formation, our experimental results show that Cl^-^ binding to GlyT1b, in the absence of substrate binding, decreases accessibility in the extracellular pathway, apparently by enhancing the formation of the Arg57-Asp460 ion pair and thereby stabilizing the outward-occluded conformation. This observation suggests that interaction of Cl^-^ with Gln299 enhances formation of the Arg57-Asp460 ion pair, independent of substrate. Our findings are inconsistent with proposals that require substrate to be bound for Cl^-^ to close the extracellular permeation pathway (36).

If the effect of Cl^-^ on extracellular pathway accessibility of GlyT1b (Fig. 4A) results from formation of the Arg57-Asp460 ion pair, then Gln299 is likely to be involved. Indeed, mutation of Gln299 prevents the conformational effect of Cl^-^ (Fig. 6B). In addition, replacement of Gln299 with glycine or asparagine also decreased Cl^-^ affinity (Fig. 6A), which could account for part, or all, of the insensitivity of these mutants to Cl^-^. Replacing Gln299 with glutamate or lysine had more dramatic effects. In Q299E, the inserted glutamate would be expected to interact strongly with Arg57 and to prevent it from forming an ion pair with Asp460. The result is a transporter locked in an outward-open conformation, unable to respond even to Na^+^ (Fig. 6B). Conversely, a lysine replacing Gln299 might be expected to interact with the bound Cl^-^ but not with Arg57, leaving the latter free to form the ion pair with Asp460. Experimentally, this resulted in a mutant adopting an inward-open conformation (Fig. 6B), unable to respond to Na^+^, which normally would close the cytoplasmic pathway.

Taken together, our results support a mechanism of Cl^-^ action on the conformation of GlyT1 and other Cl^-^-dependent NSS transporters in which Cl^-^ interaction with the conserved Gln299 allows it to disengage from Arg57, which is then free to ion-pair with Asp460, favoring the closed state of the extracellular pathway. Although not sufficient by itself, the effect of Cl^-^ binding, together with the influence of substrate binding on conformation, further stabilizes the closed extracellular pathway and facilitates opening of the cytoplasmic pathway to release substrate and ions to the cytoplasm.

## Materials and Methods

### Materials

A plasmid bearing the cDNA for N-terminal Flag-tagged human Glyt1b was a generous gift from Dr. Manuel Miranda, U. of Texas, El Paso. Monoclonal anti-Flag antibody (M1) was from Sigma-Aldrich. 2-Aminoethyl methane thiosulfonate hydrobromide (MTSEA) and [2-(trimethylammonium)ethyl] methane thiosulfonate bromide (MTSET) were purchased from Anatrace. CHIBA-3007 and desmethyl-CHIBA-3007 were generous gifts of Dr. Kenji Hashimoto, Chiba University, Japan. [^3^H]CHIBA-3007 was synthesized by methylation of its precursor, desmethyl-CHIBA-3007, with a radiochemical purity of 99% (HPLC) and a specific activity of 67 Ci/mmol by Vitrax (Placentia, CA). 2-^3^H-Glycine (42 Ci/mmol) was purchased from PerkinElmer. HEK 293 MSR cells, sulfosuccinimidyl-2-[biotinamido]ethyl-1,3-dithiopropionate) (sulfo-NHS-SS-biotin) and streptavidin agarose were from ThermoFisher Scientific. All other materials and chemicals were reagent grade obtained from commercial sources.

### Methods

#### Mutant Construction, Expression, Transport and Binding Assays

Glyt1b mutants were generated using the QuikChange site-directed mutagenesis system (Agilent). All mutants were constructed in WT or C62A background, and were confirmed by DNA sequencing.

HEK MSR cells were cultured according to the supplier’s protocol and were transfected using lipofectamine 2000 with plasmids bearing Flag-Glyt1b cDNA. Transfected cells were incubated for 40-48 h at 37°C with 5% CO_2_ prior to assays.

Glycine transport into the transfected cells was measured in 96-well plates as described previously (55). The extent of [^3^H]glycine accumulation was determined with a PerkinElmer Microbeta plate counter. The Cl^-^ dependence of glycine transport was determined using HEPES buffered saline (10 mM HEPES, 0.1 mM calcium gluconate, 1 mM MgSO_4_, 5 mM K_2_SO_4_, 10 mM glucose, 200 mM NaCl, pH 7.4) in which some or all of the Cl^-^ was replaced with gluconate to maintain a final concentration of 200 mM.

Binding of [^3^H]CHIBA-3007 was measured in cell membranes from transfected HEK MSR cells in binding buffer (10 mM HEPES buffer, pH 7.4, containing 150 mM NaCl or equimolar concentration of other salts as indicated), according to the protocol as described previously (56). To measure the effect of ions on CHIBA-3007 and glycine binding, the indicated concentrations of cold CHIBA-3007 or glycine were added to the washed membranes and pre-incubated for 5 min in binding buffer containing 150 mM NMDG gluconate, NMDG chloride, sodium gluconate, or NaCl as indicated. CHIBA-3007 binding was measured using a PerkinElmer Microbeta counter after incubation with [^3^H]CHIBA-3007 at a final concentration of 1 nM for 1.5 h at RT.

#### Cystine Accessibility Measurements

Conformational changes were measured using the accessibility of cysteine residues placed in the cytoplasmic (S244C) and extracellular (Y60C) permeation pathways. For measurement of extracellular pathway accessibility, measurements were made with intact cells expressing Glyt1b Y60C-C62A growing in 96-well culture plates. For cytoplasmic pathway accessibility, measurements were made with membranes prepared from cells expressing Glyt1b S244C-C62A on filters in 96-well filtration plates. A binding assay was used with membranes from disrupted cells because MTSET does not penetrate through intact cell membranes (57) and does not access the cytoplasmic pathway in intact cells. In both cases, accessibility was measured by the rate of cysteine reactivity with MTSET, as described previously (48). Glycine, where added, was present at 50 mM (in the absence of Cl^-^) and 0.5 mM (in NaCl). NMDG was used to replace Na^+^ and gluconate replaced Cl^-^. The MTSET concentration causing half-maximal inactivation was determined and used to calculate the rate constant for cysteine modification as described previously (58, 59). MTSET concentrations were calibrated using Ellman’s reagent (5,5’-dithiobis(2-nitrobenzoate)) (60).

#### Biotinylation and immunoblotting

Surface expression of FLAG-tagged WT Glyt1b and its mutants was determined using the membrane-impermeant biotinylation reagent sulfo-NHS-SS-biotin as described previously (61). Biotinylated Glyt1b proteins were analyzed by Western blotting using a monoclonal anti-FLAG antibody (M1) (1:1000), visualized and quantified with an IRDye 680RD goat anti-mouse IgG (Li-Cor, 1:10,000) using Li-Cor Odyssey CLx Infrared Imaging System.

#### Modeling of human GlyT1

A homology model of the glycine transporter-1 (GlyT1) was generated with MODELLER 9v22 (62) based on an outward facing structure (2.8 Å resolution, PDB ID 4XP4 (46)) of the *Drosophila melanogaster* dopamine transporter (dDAT). The alignment between the two sequences was obtained with AlignMe v1.2 in PST mode (63-65), and covered residues 30-163 and 186-495, i.e., all except a portion of extracellular loop 2, and the N- and C-termini (see Figure S1). The sequence identity between the target and the template is 46%, and thus we expect that the Cα trace of the GlyT1 model is accurate to within 1 Å of the native structure (66). Na^+^ and Cl^-^ ions in the binding site were modeled on those observed in the dDAT template structure. The top 25% of the 2000 models generated were selected according to the Modeller molecular probability density function (molpdf), and of those, the model with the highest ProQM score (67) was selected for visualization. The analysis of the homology models was performed using in-house code (https://github.com/Lucy-Forrest-Lab/hm_analysis_tool). The ProQM score for the best model is 0.842, while the average of ProQM score for the best 100 models is 0.832. These values compare favorably with the ProQM score of the dDAT template, which is 0.835. Input files and the final selected model are available on Zenodo at http://doi.org/10.5281/zenodo.4021063.

Figures of structures and models were generated using PyMol v2.3.0a0 (Schrödinger Inc.).

#### Molecular dynamics simulations of LeuT

We performed three repeats of 2 μs-long molecular dynamics simulations of the outward-open (PDB ID 3TT1, (24)), the outward-occluded (3F48, (68)) and the inward-open (3TT3, (24)) conformations of LeuT. Sodium was bound at Na1 and Na2 sites in the outward-facing states, and alanine was present in the central binding site only in the outward-occluded state. These simulations were extended from previously published ones (28, 31, 50). Briefly, each protein system was embedded in a hydrated dimyristoylphosphocholine (DMPC) bilayer using GRIFFIN (69). Sodium and chloride ions were added to achieve 100 mM NaCl concentration. The simulations of the outward- and inward-open states were performed with constant area in the *xy* plane of the bilayer, while the outward-occluded simulations were performed in the NPT ensemble. All simulations were performed using NAMD v2.9 or v2.12 (70) with the all-atom CHARMM 36 (71-74) force field. Temperature was kept constant at 310K through Langevin dynamics. The pressure was set to 1 atm with a Nosé-Hoover Langevin barostat. The non-bonded interactions were switched off smoothly from 10 to 12 Å. Long range electrostatic interactions were computed with particle mesh Ewald summations (75).

Analysis of the simulations was carried out using VMD (76). Representative simulation data are available on Zenodo at http://doi.org/10.5281/zenodo.4293733.

#### Data Analysis

Nonlinear regression fits of experimental and calculated data were performed with Origin (OriginLab, Northampton, MA), which uses the Marquardt-Levenberg nonlinear least-squares curve-fitting algorithm. The statistical analyses given were from multiple experiments and performed using Student’s paired t test.

## Acknowledgments

Acknowledgements – This research was supported in part by NIH Grant NS102277 (to G.R.), by the Guangdong Basic and Applied Basic Research Foundation (2019A1515011569) and National Natural Science Foundation of China (32071233) (to Y.-W.Z.), and by the Division of Intramural Research of the NIH, National Institute of Neurological Disorders and Stroke (to L.R.F.), and utilized the computational resources of the NIH HPC Biowulf cluster (http://hpc.nih.gov). We thank Dr. Kenji Hashimoto for his gifts of CHIBA-3007 and the desmethyl precursor. We also thank Drs. Poul Nissen and Azadeh Shahsavar for sharing their pre-publication structure of GlyT1 in an inward-open conformation.

## Author Contributions

Y-W. Z., L.R.F. and G.R. designed research; Y-W.Z., S.U., V.L., R.T.B., and N.S. performed research; Y-W.Z., L.R.F. and G.R. analyzed data and wrote the paper with input from all authors.

## SI Appendix

### Supplementary Figures

**Figure S1.**
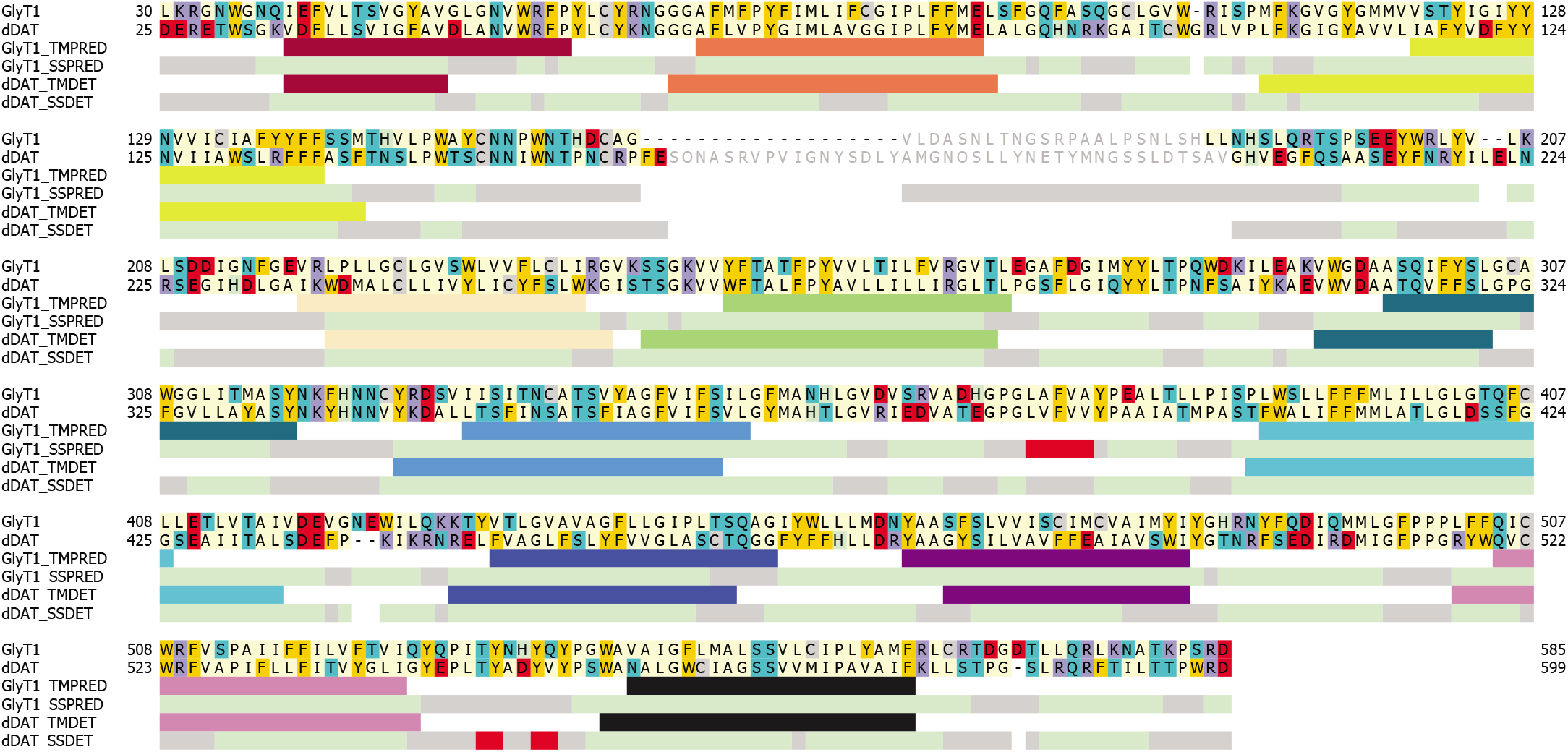
Alignment between dDAT and GlyT1 used to guide the homology modeling. Residues colored in gray are not present in either the template and not built into the model. Amino acid coloring indicates chemical properties, i.e., red for acidic (Asp, Glu), blue for basic (Arg, Lys), teal for polar (Asn, Gln, Ser, Thr), gold for aromatic (Phe, Tyr, Trp), gray for cysteine, green for histidine, and pale yellow for all hydrophobic residues, proline, and glycine. To assess the quality of the alignment we compared the transmembrane spanning regions of dDAT (“TMDET”, computed by OPM for PDB ID 4M48) (77) and GlyT1 (“TMPRED”, predicted by TOPCONS (78)). In those segments, the repeat containing TMs 1-5 is colored in red-to-green while the repeat containing TMs 6-10 is colored in blue tones. The secondary structure of dDAT (“SSDET”, computed with DSSP v3.0.1 (79, 80)) and GlyT1 (“SSPRED”, predicted with PSIPRED v4.0 (81)) is also provided: Here, the helical portion of the sequence is shown in light green, the coil regions in grey, and the extended segments in red.

**Figure S2.**
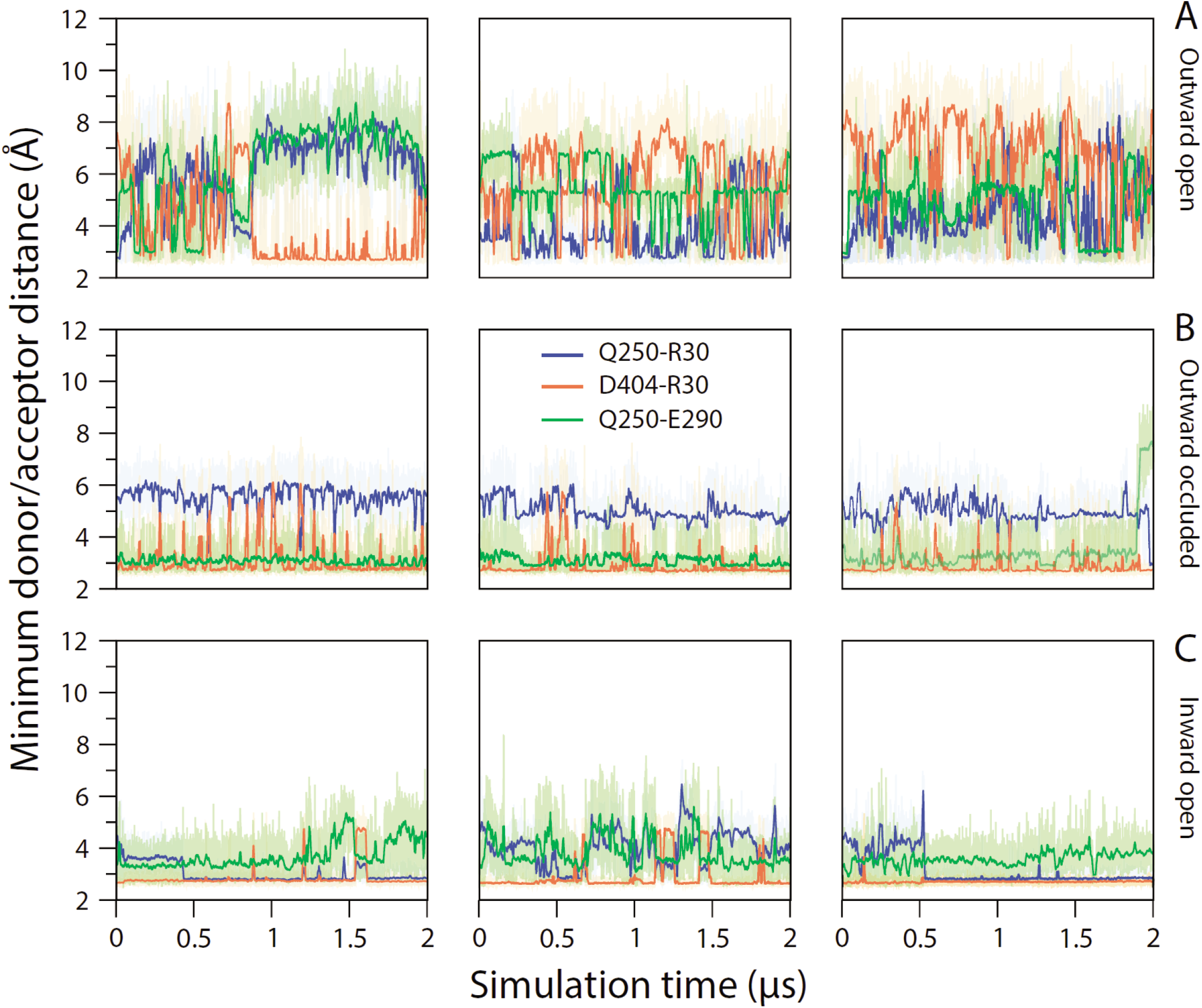
Time-course ofthe interactions between residues in the extracellular pathway and the chloride binding site during MD simulations of LeuT. Data represent distances between the closest donor and acceptor atom ofArg3O with Gln25O (blue), with Asp4O4 (orange), and between Gln25O and Glu290 (green). Simulations are for LeuT in (A) outward-open apo (3TT1), (B) outward-occluded holo (3F48), and (C) inward-open apo (3TT3) conformations. Each simulation was repeated three times (columns).

